# Integrating Engineered Macro Vessels with Self-assembled Capillaries in 3D Implantable Tissue for Promoting Vascular Integration In-vivo

**DOI:** 10.1101/2020.07.07.190900

**Authors:** Lior Debbi, Barak Zohar, Yulia Shandalov, Shulamit Levenberg

## Abstract

Fabrication of a functional hierarchical vascular network remains an unmet need for cultivation and transplantation of 3D engineered tissues. In this work, an effective approach was developed to fabricate a functional, perfusable and biocompatible, multi-scale vascular network (MSVT) within thick, implantable engineered tissues. Using a templating technique, macro-vessels were patterned in a 3D biodegradable polymeric scaffold seeded with endothelial and support cells within a collagen gel. The lumen of the macro-vessel was lined with endothelial cells, which further sprouted and anastomosed with the surrounding self-assembled capillaries. Anastomoses between the two-scaled vascular systems displayed tightly bonded cell junctions, as indicated by vascular endothelial cadherin expression. Moreover, MSVT functionality and patency were demonstrated by dextran passage through the interconnected hierarchical vasculature. Additionally, physiological flow conditions were applied with home-designed flow bioreactors, to achieve a MSVT with a natural endothelium structure. Finally, implantation of a multi-scale-vascularized graft in a mouse model resulted in a clear beneficial effect, as reflected by extensive host vessel penetration into the graft and an increase in blood perfusion via the engineered vessels as compared to control microscale-vascularized graft. Designing and fabricating such multi-scale vascular architectures within 3D engineered tissues is essential, both for in-vitro models and for therapeutic translation research.

## Main Text

### Introduction

One elementary need of real-scale engineered tissues with potential to repair damaged organs is a sufficient blood supply providing proper exchange of nutrients and gasses essential for cell survival. Many attempts to engineer vascularized tissue have been made over the last two decades. Some of the common approaches aim to artificially fabricate channel networks embedded within the engineered tissue by means of direct bioprinting of vessel-like structures(1–3) or indirect printing of 3D channel templates using soluble sacrificial materials, followed by endothelial cells (ECs) lining(4–7). Yet, these strategies have shown limited success as they fail to meet vascular network hierarchy and complexity and could not maintain channel network structure under internal contractile forces exerted by supportive cells. Additionally, most of the solutions rely on soft hydrogel-based scaffolds that are difficult to handle for transplantation.

Alternatively, the inherent ability of ECs to spontaneously self-assemble into living capillaries in the presence of supportive cells can be harnessed to establish well-structured micro-vascular networks (8, 9). The engineered microvasculature has been shown to integrate and function upon implantation (10, 11). However, the self-assembled micro-vessels are not organized in a hierarchical structure, consequently limiting the scale, maturation and integration level of the engineered tissue.

Several attempts have been made to fabricate a perfusable, hierarchical vessel network in a dish, mainly for micro lab-on-a-chip systems. This includes the works of Chen group, MIMETAS Ltd. and others, in which engineered vessels were fabricated within natural hydrogels and sprouting angiogenesis of capillaries was stimulated by pro-angiogenic factors (12–14). Yet, these endothelial-based systems are limited in their ability to reconstruct 3D functional engineered tissue which requires multiple cell combinations and mechanical durability for handling. Several groups, including Kamm group, Hughes group and others were able to show multi-scale vascular structure in co-culture on a chip (15–18); however, the micro-chip systems are not suitable for implantation, as they are neither made of biodegradable material nor on a tissue scale.

An additional approach relies on the micro-fabrication of channel networks within a synthetic chip using a molding technique which enables perfusion through the channels (19, 20). However, the synthetic nature of the channel network precludes the desired biological integration of the chip with the host.

To address the unmet need for in vitro-fabricated, functional, multi-scale vascular network within thick, implantable grafts, we developed a novel approach that establishes multi-scale vascular networks (MSVTs) comprised of patterned macro-channels integrated with self-assembled micro-vessels, within 3D engineered tissues. The vascularized engineered tissue is formed within a biodegradable, highly porous, polymeric scaffold, which provides mechanical support for the tissue structure to withstand internal cellular contractile forces and external forces that evolve during the cultivation period, handling and implantation in-vivo. We also studied the effect of various early and late wall shear stress stimuli levels on endothelium structure using a compatible pump-free perfusion bioreactor, and defined the flow patterns required for the MSVT function. Finally, we implanted the MSVT tissue graft in a mouse model to assess its effect on vessel integration and graft perfusion.

## Results

### Construction of multi-scale vasculature in an implantable engineered tissue

We developed a strategy that enables the integration of artificially patterned macro-vessels (500-1000μm) with naturally self-assembled capillaries (10-80μm) in an implantable platform. 3D printing was used to establish a polydimethylsiloxane (PDMS) mold that can later be removed before implantation, and with precise placements for templating rods (Figure 1Ai). A salt-leached porous scaffold made of a poly(l-lactic acid)/poly(lactidecoglycolide) (PLLA/PLGA) co-polymer was cast inside the PDMS mold to mechanically support the engineered tissue (Figure 1Aii). Thereafter, a cell mixture of human adipose microvascular endothelial cells (HAMEC - tdTomato) and dental pulp stem cells (DPSC) were seeded within collagen gel to form self-assembled capillaries (Figure 1Aiii). After gel solidification, the two templating rods were removed and the channels were uniformly seeded with endothelial cells (ECs, HAMEC - ZSgreen), to form an endothelial lumen within the macro-vessels (Figure 1Aiv & 2A,B). During cultivation, ECs located in the lumen sprouted and anastomosed with the peripheral self-assembled capillaries, to form the MSVT (Figure 1Av & B). On day 1, peripheral co-cultured ECs rearranged into small clusters and spontaneously formed a branched vascular network by day 4 (Figure 2C). The self-assembled micro-vessels reached a total length of 31.6±4.3mm by day 4, which slightly increased by day 7 (36.3±5.1mm). The average length of the micro-vessels reached 117.2±15.8μm by day 4 and remained stable until day 7 (Figure 2C). Moreover, the MSVT was associated with significantly higher vascular network development in the depth of the tissue when compared to one scale self-assembled vasculature (Figure S1). ECs sprouting from the macro-vessels showed various integration patterns with the peripheral self-assembled micro-vessels, forming wrapping, mosaic and end-to-end vessel interconnections (Figure S2). The connectivity of the anastomosed micro-vessels was demonstrated by the integrity of VE-cadherin between both types of labeled ECs. (Figure 2D).

**Figure 1.**
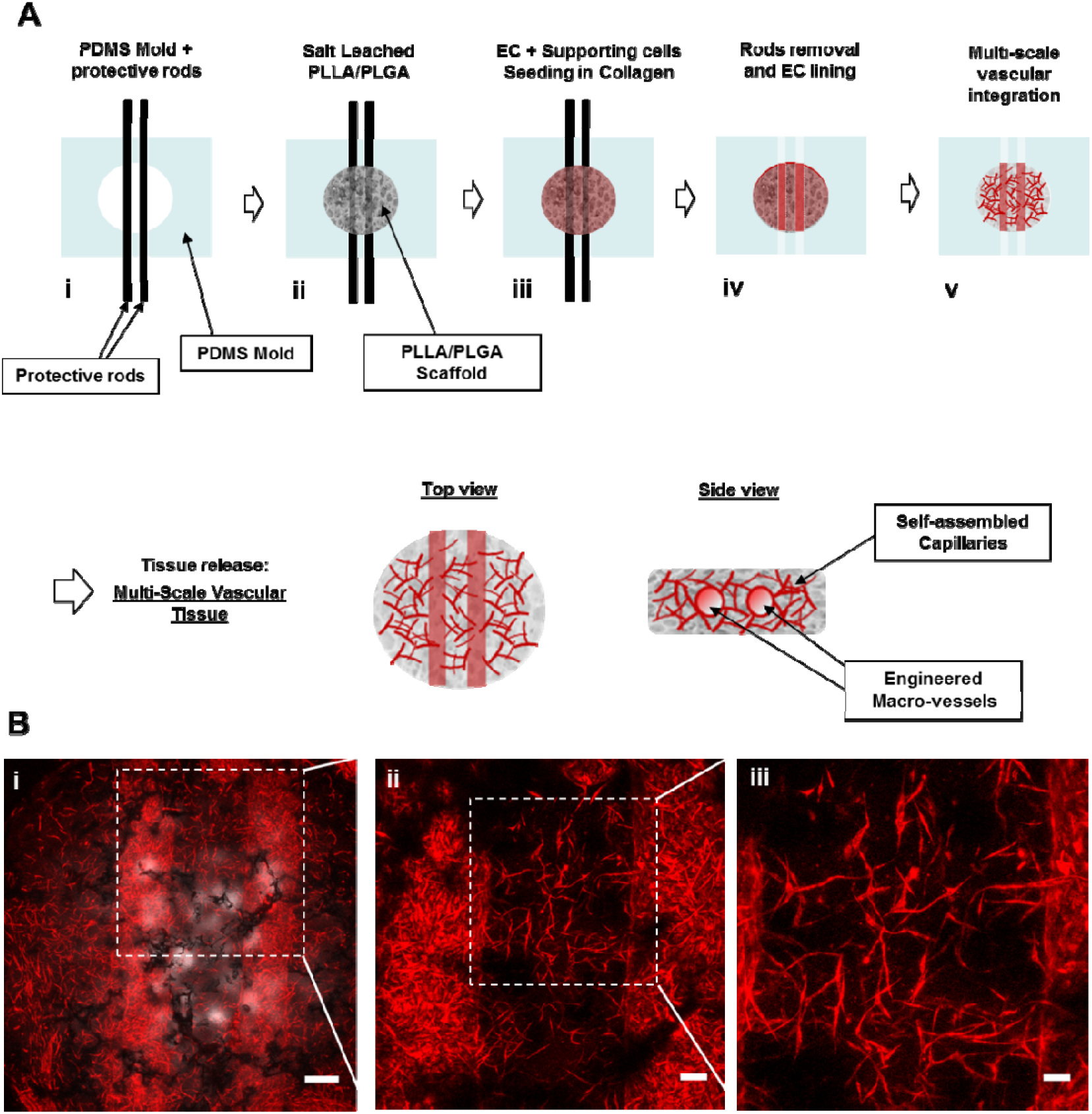
Multi-scale vascular tissue (MSVT) construction. (A) Schematic of the MSVT fabrication process. (i) Macro-vessel templating rods are placed within a supporting PDMS house. (ii) Preparation of a porous PLLA/PLGA scaffold using a salt-leaching method. (iii) Casting collagen gel seeded with ECs and SCs. (iv) Fabricating engineered macro vessels by templating rods removal followed by ECs lining (v) Construct in-vitro cultivation forming multi-scale vascular tissue. (B) Maximum intensity projection of the MSVT, with high magnifications. HAMEC tdTomato (red) co-cultured with DPSCs (not labeled). The two engineered macro vessels anastomose with the surrounding micro-vasculature can be observed. The PLLA/PLGA scaffold can be indicated in (B1) image via bright-field channel (black shadow indicating the scaffold). (Scale bar: 500μm (left), 100μm (middle), 50m (right))

**Figure 2.**
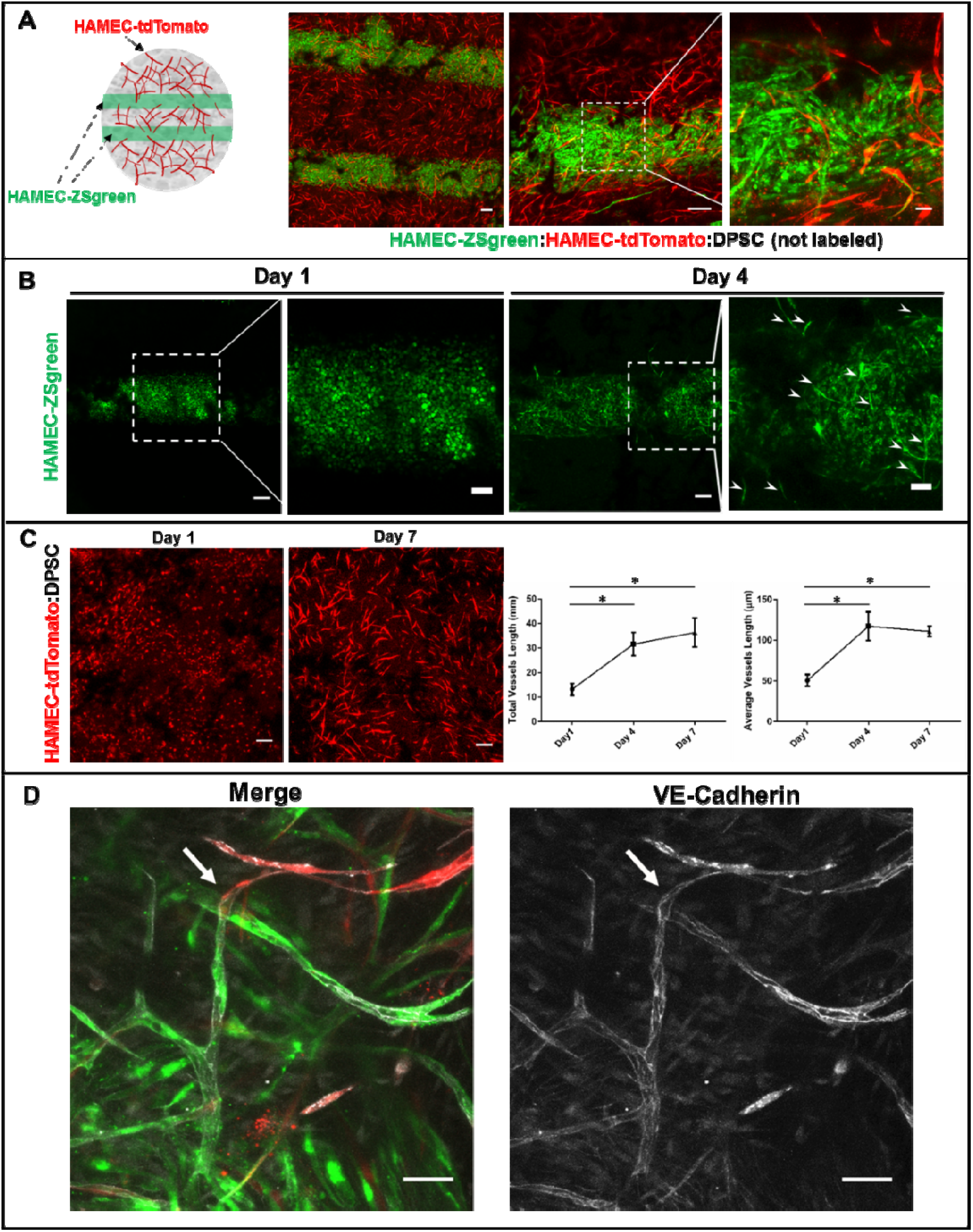
Multi-scale vascular tissue development. (A) Schematic and several magnifications of representative MSVT images. Macro engineered vessels lined with HAMEC ZSgreen (green) surrounded by HAMEC tdTomato self-assembled micro vasculature (red) co-cultured with DPSCs (not labeled). (Scale bar: 100μm (left), 100μm (middle), 50m (right)) (B) Sprouts emerging from a macro vessel, as captured 4 days post seeding. (Scale bar: 200μm (zoom-out), 50μm (zoom-in)) (C) Micro vasculature development over time, including quantification of total and average vessel length (measured at days 1,4,7). (Scale bar: 200μm) (D) VE-cadherin staining (white), indicating functional interconnected green sprout emerging from engineered a macro-vessel with a red self-assembled capillary. (Scale bar: 50μm) Data are presented as means ± SEM. N ≥ 4 (*P < 0.05).

### Multi-scale vasculature connectivity and patency

To investigate the ability of the multi-scale vasculature to distribute blood from macro-vessels to capillaries, fluorescently labeled 10kDa dextran was injected into the patterned macro-vessels. The dextran was captured inside the macro-vessels due to the size-selective endothelial barrier (Figure 3A), which prevented spontaneous outer diffusion. Rather, we inspected dextran inside sprouted luminal ECs (Figure 3B). These sprouted luminal ECs resulted in anastomoses with self-assembled capillaries (Figure 3C) which exclusively enabled the transfer of dextran from the patterned macro-vessel into the open self-assembled vascular network (Figure 3D).

**Figure 3.**
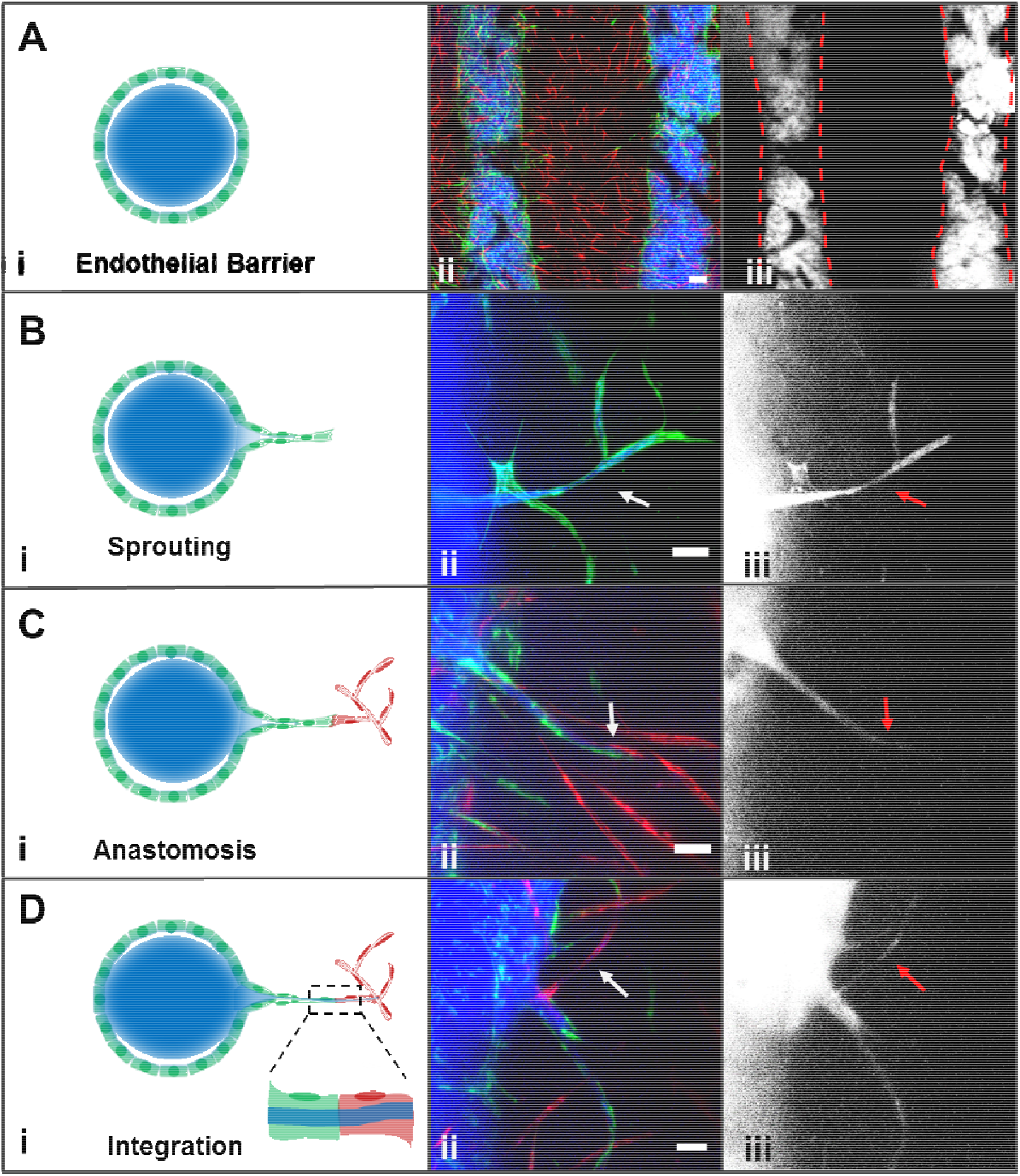
Multi-scale vascular tissue integration and patency. Cascade blue fluorescent dextran (blue) was injected into macro vessels lined with HAMEC ZSgreen (green) anastomosed with surrounding microvasculature formed by HAMEC tdTomato (red). (A) Macro vessels endothelium serves as a barrier and holds injected 10kDa dextran. (B) Open functional sprout emerged from engineered macro vessel as can be indicated by dextran entrance to the hierarchical structure. (C) Sprout anastomosis with the microvasculature. (D) Complete integration of a sprout with the microvasculature results with open functional lumen and dextran entrance. (Scale bar: 200μm (A), 20μm(B-D))

### The impact of flow on the multi-scaled vascular network

In order to study the influence of flow on MSVT, both on endothelium structure and sprouts emergent, we designed and fabricated a pump-free flow bioreactor to enable the application of luminal flow inside the macro-vessels (Figure S3). The bioreactor was designed to generate luminal flow conditions inside the macro-vessels by continuously altering the hydraulic head using an external tilting plate (Figure 4A). In these experiments we tested the impact of different levels of physiological shear stresses, applied at early and late stages of culture under static conditions (Figure 4B) on endothelium arrangement and luminal ECs sprouting. We hypothesized that flow-induced shear stress within physiological range will affect endothelium structure and sprouts emergent. Hence, we tested the impact of flow before (early induction) and after sprouting emerged (late induction). First, we explored the impact of flow-induced wall shear stimuli of 0 (static), 1 (low), and 10 dyne/cm^2 (high). Under static conditions, the morphology of the luminal ECs within the macro-vessels developed into spindle like elongated structure (Figure 4C-E). On the contrary, both low and high shear stress levels induced changes in EC morphology indicating on cell arrangement into endothelium structure (Figure 4C and S4). Under flow conditions, luminal ECs became more rounded, less dense and formed a spacious organized EC layer compared to static conditions (Figure 4C and S4). To quantify the change in EC morphology, we measured and compared the mean cell aspect ratio. Both early and late flow protocols significantly decreased luminal EC mean aspect ratios, from 4.3±0.47 and 3.3±0.19, respectively, to values close to 1 (1.34.3±0.05 and 1.5±0.03, respectively; Figure 4E), reaching a typical cobblestone endothelium structure. Additionally, we assessed the arrangement of ECs into endothelium by staining VE-cadherin which is indicative intercellular junction. Under flow conditions, VE-cadherin patterns revealed the natural and typical cobblestone-like structure of endothelium, while under static conditions, elongated spindle-like ECs structure was observed (Figure 4C).

**Figure 4.**
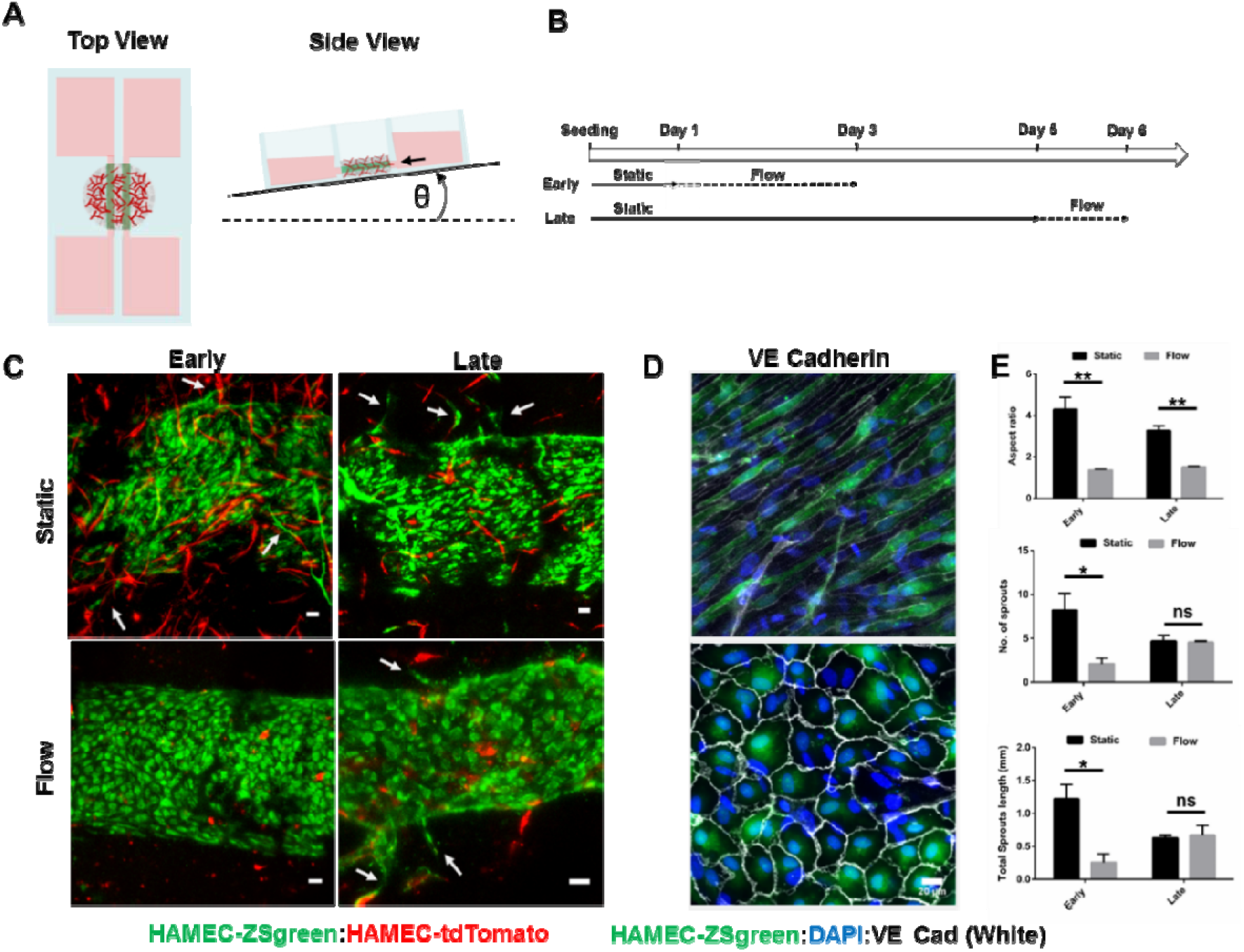
Establishing an endothelial structure under flow conditions. (A) Schematic of a pumpless flow bioreactor. Flow is induced using a tilting platform. (B) Timeline of early and late flow induction profiles within the MSVT. (C) Maximum intensity projection images of macro vessel endothelium and sprouts under static or flow conditions for early and late profiles. (Scale bar: 50μm) (D) VE-cadherin (white) and DAPI (blue) staining of a HAMEC ZSgreen macro vessel endothelium under static (top) and flow (bottom) conditions. (Scale bar: 20μm) (E) Quantification of endothelium cell aspect ratios, sprout number and sprouts length under static and flow conditions both for early and late flow profiles. Data are presented as means ± SEM. N ≥ 3 (*P < 0.05, **P < 0.01).

Furthermore, under static conditions we observed EC sprouting initiated from the macro-vessel lumen (Figure 4C). Throughout this process, ECs invaded the surrounding engineered tissue and sprouts continued to elongate until they anastomosed with self-assembled micro-vessels (Figure 2A, 2D and S2). In agreement to former studies(21, 22), early flow conditions resulted in a significant decrease in luminal EC sprouting, as measured by both the number and total length of sprouts (Figure 4C,E). However, the application of flow at a later stage, after sprout emergence (day 5), resulted with typical cobblestone endothelium structure, while sprout number and length were maintained as compared to static conditions (Figure 4C,E).

### MSVT graft integration upon implantation

To investigate the capability of MSVT to integrate with host vasculature in vivo, we compared between host vascular integration with MSVT and micro-scale vascularized tissues (VT). For this end, MSVT and VT were cultivated in vitro until reaching fully developed vasculature pre-implantation. Then, constructs were released from the PDMS molds without damaging the vascularized tissue (Figure S5) and implanted subcutaneously in the same nude mouse (Figure 5A).

**Figure 5.**
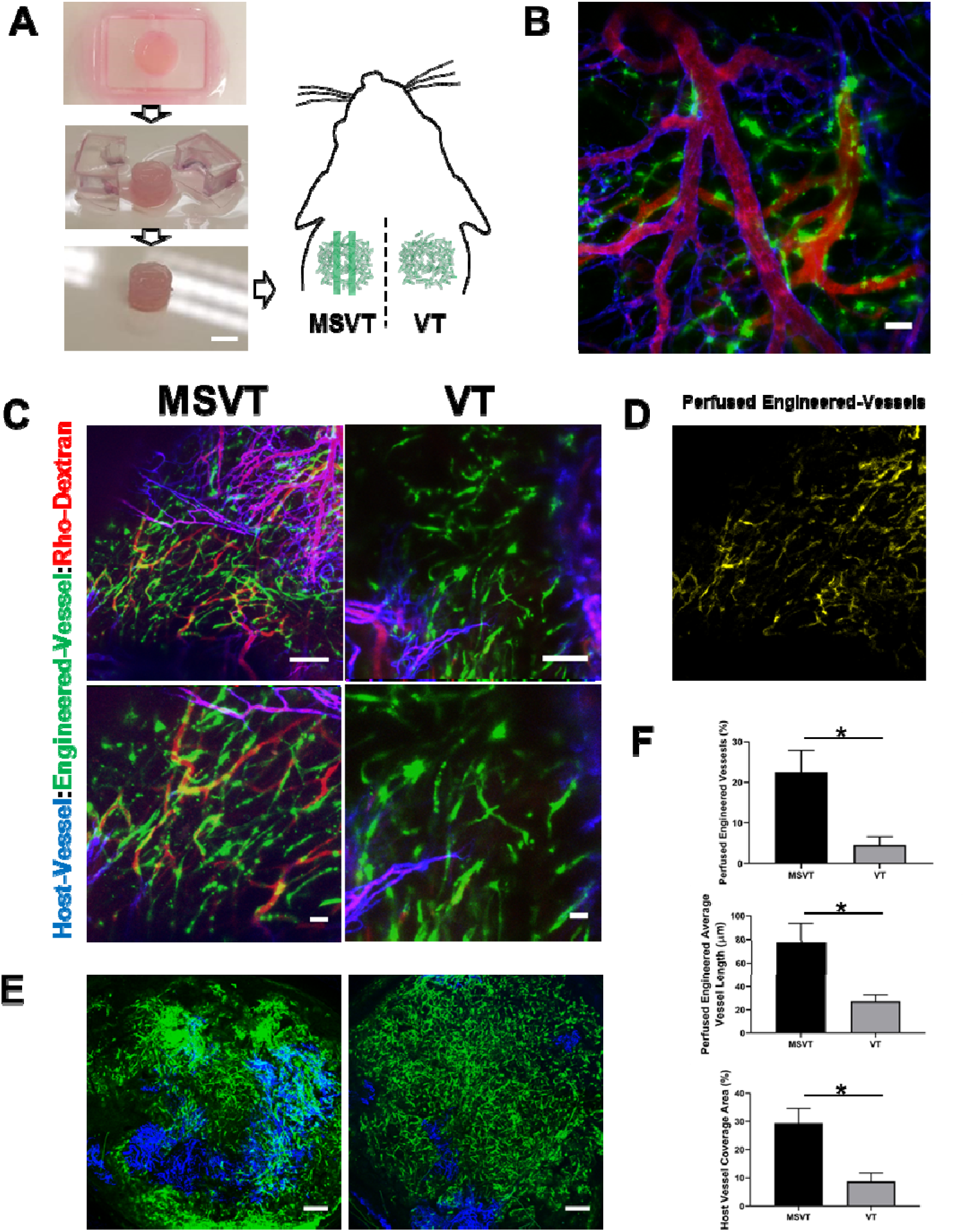
Multi-scale vascular tissue (MSVT) promotes in vivo vascular integration. (A) Images of construct release before implantation (left) and schematic subcutaneous MSVT and VT implantation in a mouse model (right). (Scale bar: 5mm) (B) Representative image of in-vivo MSVT integration and perfusion of engineered vessels (green) and host vessels (blue) with TRITC-dextran (red). (Scale bar: 50μm) (C) Representative images of in-vivo perfused vasculature for MSVT and VT. (Scale bar: 200μm (upper), 50μm (lower)) (D) Co-localization of dextran and perfused engineered vessels in MSVT. (E) Representative images of host vessel invasion (blue) into MSVT and VT constructs. (Scale bar: 500μm) (F) Comparison between implanted MSVT and VT with regards to perfused engineered vessel percentage, their average length and host vessel invasion into graft 8 days after implantation. Data are presented as means ± SEM. n = 4 (*P < 0.05).

After 8 days, graft-host vascular integration was assessed by staining the host vessels with Alexa-647-anti-CD31 antibody and tracing the blood perfusion in the graft by injecting tetramethyl-rhodamine-isothiocyanate (TRITC)-dextran intravenously. Further, the grafts were immediately dissected and scanned by confocal microscope. The engineered micro-vessels were maintained complete and visible both in the MSVT and VT. However, the MSVT resulted in an enhanced graft-host vascular integration indicated by more vessel anastomoses and by the existence of dextran inside the engineered vessels (Figure 5B,C). We measured the percentage of perfused engineered vessels (e.g. the total engineered vessels that were perfused with dextran out of the total engineered vessels (Figure S6). The MSVT showed significantly enhanced integration and functionality with the host vasculature as compared to the VT graft, as reflected by the five-fold higher rate of total perfused engineered vessels (22.5±5.3% in MSVT, 4.55±2.1% in VT grafts, Figure 5C, F). Furthermore, the perfused vessels in the MSVTs showed a significantly higher average vessel length as compare to the VTs (77.9±15.5μm versus 27.4±5.5 μm, respectively, Figure 5F). In addition, the MSVTs revealed significantly higher host vessel invasion into the grafts and large implant coverage (29.37±4.5%) as compared to the VT (8.77±2.6%) as can be observed in Figure 5E, F.

## Discussion

Engineered tissues designed for clinical applications must contain a functional vascular network to support blood supply upon implantation. Naturally, a hierarchical feeding vasculature including multi-scale vessels is essential to support highly cell-dense tissues and organs. For example, blood vessels feeding the heart, pancreas and liver tissues include arteries and veins in the scale of 1mm, connected to capillary networks of micrometric scale (23). While multiple studies have focused on either macroscale (1–5, 24) or microscale vasculature fabrication (9, 11, 25, 26), this study combined patterned macro-vessels (600μm) and self-assembled micro-vessels (10-80μm) within a single system, to generate a multi-scale vasculature, relevant for therapeutic applications.

One challenge in fabricating MSVT is the ability to maintain the vascular structure while exposed to cellular contractile forces during in-vitro cultivation. It was shown that cells seeded within soft hydrogel-based scaffolds (natural or synthetic) exert contractile forces which induce large deformations and construct shrinkage overtime (27). To overcome this critical challenge, the MSVTs in the presented system were mechanically reinforced with a biodegradable polymeric (PLLA-PLGA) scaffold. This salt-leached polymeric scaffold is highly porous (~93%), mechanically durable, highly biocompatible, and can be degraded post-implantation in a tunable way (26, 28, 29). Embedment of this supportive polyester scaffold in the MSVT, maintained the construct shape, including the multi-scale vascular network architectures, and prevented shrinkage during the in-vitro cultivation period (Figure 1B). Furthermore, the supportive polymeric scaffold enabled tissue handling, including application of external mechanical loads required for tissue release from the PDMS mold and implantation (Figure S5).

The micro-scale vasculature in the MSVT formed upon the spontaneous arrangement of ECs into capillary networks after 3-4 days of co-culture with supportive DPSCs (Figure 2C) (30). The natural EC self-assembly process offers intrinsic advantages over artificial micro-vessel fabrication techniques, such as laser-guided patterning, droplet printing and micro-groove templating (31–36). Self-assembled vasculature is naturally driven, doesn’t require complex design and fabrication techniques, forms in optimal scale, maintains high cell viability, dynamic and can change according to the tissue needs via angiogenesis process. In the MSVT, the ECs lining the macro-vessel lumen responded to pro-angiogenic factors secreted by the tissue and sprouted towards the surrounding self-assembled vasculature (Figure 1A, B). The sprouted luminal ECs fully anastomosed with the self-assembled vasculature, as indicated by VE-cadherin staining (Figure 2D). Interestingly, luminal EC sprouts integrated with the self-assembled vasculature via three different types of vessels interconnections, i.e., vessel wrapping, host-graft vessel mosaic and end-to-end vessels interconnections (Figure S2), as has been previously reported in in vivo studies (11, 37).These findings demonstrate the ability of the MSVT to mimic natural biological mechanisms essential for vascular integration, in vitro.

The presented MSVT contains engineered macro vessels lined with ECs which form a uniform functional endothelium. As was shown in another work, engineered macro vessels are characterized by a selective barrier for large molecules(22). Dextran 10kDa represent such molecules circulated in the blood. In this work, we demonstrated the feasibility of transferring 10kD dextran, blocked by the endothelial barrier (Figure 3A), from macro vessel into self-assembled microvasculature in an implantable 3D tissue (Figure 3B-D). This finding indicates the connectivity and patency of the MSVT.

The influence of shear stress on macro vessels EC morphology and behavior was studied previously within a cell-free collagen matrix(22, 38). Here, we studied its effect on endothelium EC behavior within a 3D highly cell-dense engineered tissue. Using our in-house-designed flow bioreactor (Figure S3), we tested the effect of static conditions, low (1 dyne/cm^2) and high (10 dyne/cm^2) shear stress levels which were chosen to fit physiological range(23, 39). In agreement with other studies(40, 41), under flow, the macro vessel endothelium morphology developed into typical cobblestone-like structure built from rounded luminal ECs with low aspect ratio values (Figure 4C-E, Figure S4), while under static conditions, luminal ECs formed elongated shape accompanied with high aspect ratio values (Figure 4C-E) indicating on higher cell motility(42) triggered by pro-angiogenic conditions, which leads to sprout initiation. The effect of shear stress on sprouting angiogenesis in engineered macro-vessel models is still under debate. While some studies have shown sprouting inhibition and quiescence upon shear stress stimuli (21, 22), others report on enhanced sprout growth(38, 43). In the presented MVST we observed inhibition effect on new sprouts emergent from the macro vessel endothelium under early shear stress application (1-day post-seeding, Figure 4C,E), while static conditions lead to a significant emergence of new sprouts from the macro-vessel (Figure 4C,E). Thus, to obtain both sprouts emergent and typical cobblestone-like endothelium structure, we developed a two-stage protocol. At first, cells are cultured under static conditions for 5 days, to establish the conditions for angiogenesis. At this stage, luminal ECs sprout and anastomose with the self-assembled vasculature, likely driven by the pro-angiogenic gradient which develops over time. The second step includes the application of physiological wall shear stress for a relatively short duration (24 h), which triggers the formation of the typical endothelium structure. More specifically, the EC sprouts are maintained, while the luminal EC aspect ratio is decreased and the cobblestone endothelium structure evolves (Figure 4C, E).

In vivo implantation of engineered tissues with a mature micro-vasculature has been shown to improve graft-host integration and perfusion (11, 44, 45). However, in order to fabricate large-scale tissues, perfusable macro-vessels must be included within the engineered tissues. As shown in Figure 1S, in-vitro cultivation of the MSVT resulted in a more developed vasculature within the tissue depth as compared to VT. This can be explained by enhanced diffusion through the macro-channels, which may improve material transport to the depth of the scaffold. Subsequently, the significantly superior vascular integration in vivo (graft perfusion, vessel penetration and anastomosis, Figure 5C-F) of MSVT as compared to VT can be explained by the improved microvasculature developed in vitro at the depth of the MVST. In addition, the engineered macro-vessels possible contributed to increased host vasculature invasion by creating an open guiding paths. The successful integration of the MSVT graft lies in agreement with a previous work of our group studying the graft-host vascular integration of a vascularized engineered tissue (VT) (11). While the two studies showed similar percentages of perfused engineered vessels and host vessel penetration, the MSVT in this study had significantly shorter cultivation time (6 days vs. 14 days) and shorter implantation time in-vivo (8 days vs. 14 days). Moreover, when comparing the results for similar in-vitro and in-vivo time frames as reported in the compared study (7 days in-vitro, 10 days in-vivo), the MSVT showed 3-times higher values for the analyzed integration parameters, emphasizing the clear superiority of the hierarchical multi-scale vascular network over micro vasculature within engineered tissues upon implantation.

Multiple studies have shown that the self-assembled microvasculature forming within 2.5D micro-chip assays can anastomose successfully to a perfusable engineered macro-vessel (15, 16, 25). While these studies reported on perfusable hierarchical vasculature only within micro-chip assays (100μm thick), we managed to form multi-scale interconnected vascular network in a real 3D engineered tissue of large-scale dimensions (7mm diameter and 2mm thick), relevant for graft implantation. The interconnectivity and patency demonstrated in the multi-scale vascular network is important for enabling long-term tissue feeding in-vitro. By allowing transport into self-assembled vasculature, the MSVT will enable to study the interaction between perfused functioning micro vessels and the surrounding tissue (e.g. vascular diseases models, cancer metastasis etc.) in a more physiological model. Furthermore, in contrast to photolithography-based micro-fabrication, the presented approach can be easily modified and scaled up to engineered tissues of therapeutically relevant thickness, by combining 3D printing technology to fabricate the macro-vessel network and introducing endothelial cells within the surrounding printed bioink that will organize spontaneously into micro vessel network. In conclusion, EC-lined macro-vessels can anastomose with self-assembled micro-vessels to create a hierarchical multi-scale vascular 3D tissue. This hierarchical architecture has a beneficial effect upon implantation by promoting rapid vasculature integration and graft perfusion, which are essential for achieving high viability and functionality of large engineered tissues post-implantation.

## Materials and Methods

### Cell culture

Human adipose microvascular endothelial cells expressing green fluorescent protein (HAMEC ZS green) and HAMEC expressing red fluorescent protein (HAMEC tdTomato) (ScienceCell) were cultured in endothelial cell medium (ScienceCell, Cat. No. 1001). Dental pulp stem cells (DPSCs) (Lonza, USA) were grown in low-glucose Dulbecco’s modified Eagle medium (DMEM; Gibco), supplemented with 10% fetal bovine serum (FBS; Hyclone), 1% non-essential amino acids (NEAAs, Gibco), 1% GlutaMAX (Gibco), and 1% penicillin-streptomycin-nystatin solution (Biological Industries). All cell types were grown to passage 6-8 in T-flasks incubated in 5% CO_2_ and 37°C.

### MSVT scaffold preparation

The disc-shaped (2-mm-thick and 7mm-diameter) pattern for the MSVT was designed using SOLIDWORKS 2017 CAD software, with precise leading channels designed for placing two templating rods (parallel orientation with 1mm relative distance). Negative pattern molds were made of water-soluble butenediol vinyl alcohol co-polymer (BVOH) and printed with a PRUSA MK2.5 3D printer. The mold was cast with poly(dimethylsiloxane) (PDMS;1:10; Sylgard 184; Dow-Corning), cured for 2h, at 80°C, and washed with distilled water, overnight, to remove sacrificial material. Further, templating rods were placed and porous sponges composed of 50% poly-L-lactic acid (PLLA) (Polysciences, Warrington) and 50% poly(lactic-co-glycolic acid (PLGA) (Boehringer Ingelheim) were fabricated utilizing a particle-leaching technique, to achieve pore sizes of 212−600μm and 93% porosity, as described in previous works (26, 29).

### MSVT seeding

Pre-seeding preparation included fibronectin coating of the PLLA/PLGA scaffolds (100 μg mL^−1^, Sigma), for 1h, at 37°C, in a tissue culture incubator. HAMECs (1.3×10^5^ cells) and DPSCs (3.5×10^5^ cells) were mixed in 100μl collagen (3.5mg/ml, rat tail collagen type 1, Corning, prepared according to the manufacturer protocol). The cell mixture was then seeded onto the scaffolds and allowed to solidify for 30 min, at 37°C, 5% CO_2_. Macrovessel fabrication included gentle removal of templating rods and injection of HAMEC (1×10^7^ cells/ml) into the lumen. Uniform cell attachment to the lumen was achieved by flipping the MSVT every 10 min during 30 min incubation, at 37°C. Detached cells were washed away with HAMEC medium and samples were then incubated in a 1:1 mixture of HAMEC:DPSC culture media (8 ml per well). MSVTs were cultured for 6-12 days and medium was replaced every 2-3 days.

### Flow bioreactor design, fabrication and set up

The flow bioreactor was designed and fabricated using a methodology similar to that described for the MSVT preparation. The bioreactor was designed to enable fabrication of the MSVT scaffold in the center of the device, with 4 reservoir wells connected to the macro-vessels (Figure S3). The chambers were designed to enable media flow through each macro-vessel separately. The luminal flow was applied through each macro-vessel by changing the hydraulic heads in the reservoir using a tilting plate. The tilting angle was set to the minimum (7°) and frequencies were set to 1RPM and 10RPM to generate a mean wall shear stress of 1 and 10 dyne/cm^2^, respectively, according to the following Hagen-Poiseuille equation:

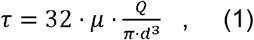

where μ is the kinetic viscosity, Q is the mean volumetric flow rate, and d the macro-vessel diameter. The Haagen-Poisseuille equation was applied under the following assumptions: the medium is considered a Newtonian fluid, the macro-vessel cross-sectional area is cylindrical, the macro-vessel is straight with inelastic walls, and the medium flow is steady and laminar. The Haagen-Poisseuille equation indicates that shear stress is directly proportional to blood flow rate and inversely proportional to vessel diameter. Mean volumetric flow rate (Q) was calculated as follows:

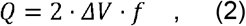

where *ΔV* is the change in medium volume measured empirically in a well before and after half a cycle for a given frequency, and f is the tilting frequency.

### Quantification of vessel network formation and development

Confocal images were acquired on days 1, 4, 7 post-seeding, using a Zeiss LSM700 confocal microscope (Carl Zeiss). To quantify total and average vessel length during vasculature formation, images were processed by ImageJ (NIH) and analyzed using AngioTool (National Cancer Institute) software.

### Whole-mount immunofluorescence staining

Scaffolds were fixed with 4% paraformaldehyde (PFA; Electron Microscopy Sciences), for 20 min, rinsed with phosphate buffered saline (PBS) (Gibco® Life Technologies) for 5 min x3 times, and permeabilized with 0.3% Triton X-100 (Bio Lab Ltd.), for 10 min at room temperature (RT). Following PBS washes (5 min x3) and overnight incubation in blocking buffer (5% bovine serum albumin (Merck Millipore) in PBS) at 4°C, scaffolds were incubated overnight, at 4°C, with goat anti-VE-cadherin antibodies (Sigma-Aldrich) diluted 1:100 in blocking buffer. Following PBS washes, samples were incubated with secondary antibodies (1:400 donkey anti-mouse Alexa Fluor®647 or 1:100 donkey anti-goat Alexa Fluor®647; Jackson ImmunoResearch) and 1:1000 4’,6-diamidino-2-phenylindole (DAPI, Sigma-Aldrich), for 3h, with gentle shaking, at room temperature. Following PBS washes (5 min, x3), scaffolds were imaged with a confocal microscope, using the Zen software. Image processing and further analyses were performed using FIJI software.

### Endothelial cell aspect ratio evaluation

Confocal images of the engineered macro-vessel endothelium were analyzed using ImageJ software. The influence of different shear stress conditions on cell shape was evaluated by manually measuring and quantifying the cell aspect ratio using A/R definition as the ratio between the cell’s long versus short axis, with 1 indicating a pure circular shape and higher values directly correlating with cell elongation.

### Evaluation of MSVT in-vitro patency

To evaluate the patency of the MSVT in-vitro, 10 mg/ml cascade-blue–dextran (average MW 10,000, Sigma–Aldrich) was directly injected into the open ends of the engineered macro vessels. The fluorescent dye was then viewed via confocal microscope imaging.

### In-vivo graft implantation

All surgical procedures and animal studies were approved by the animal ethics committee at the Technion. Athymic female nude mice (~25 g, 7-9-weeks-old; Harlan Laboratories) were anesthetized via intra-peritoneal injection of a mixture of 100mg/kg ketamine and 10 mg/kg xylazine. After 6 days of in vitro cultivation, the MSVTs and VTs were subcutaneously implanted into the back region and the incisions were closed with 4-0 silk sutures. One week post-implantation, Alexa-flour647 (AF647) anti-mouse CD31 antibodies (Biolegend) was intravenously injected into the mouse tail vein and allowed to circulate for 30 min. Mice were than anesthetized and 10 mg/ml TRITC–dextran (average MW 155,000, Sigma– Aldrich) was intravenously injected through the tail vein. Immediately after, mice were euthanized with CO_2_, grafts were retrieved, fixed in 10% formalin (Sigma-Aldrich), and scanned using a LSM700 confocal microscope.

### Evaluation of perfusion rates in the graft

The perfusion rates in the engineered vessels of the MSVT and VT grafts were quantified using AngioTool software (National Cancer Institute), using the following methodology. First, the total and average length of engineered vessels was segmented and analyzed. Next, the images of the dextran channel and the engineered vessels were intersected to create a perfused engineered vessel channel. This channel was further processed using ImageJ and segmented by the AngioTool to quantify the total and averaged perfused vessel length. The rate of perfused vessels was calculated as the percentage of segmented perfused engineered vessels out of the total number of segmented engineered vessels (Figure S6).

### Evaluation of host vessel invasion into the graft

Graft perimeter in each image was determined using the bright-field channel. Using ImageJ software, host vessels presence, marked by Alexa-flour647 (AF647)-conjugated anti-mouse CD31 antibodies (Biolegend), was evaluated by calculating the coverage area (%) of the binarized host vessel image within the graft region.

### Statistical analysis

Data are presented as mean ± SEM. Group differences were determined using the unpaired, two-tailed Student’s t test. p<0.05 was taken to indicate a statistically significant difference between groups. Statistical analyses were performed using Prism (GraphPad) Software.

## Acknowledgments

We thank Janette Zavin for assistance with the staining, Mark Tendler and Shira Landau for assistance with the animal model, and Yehudit Posen for editorial assistance in preparing this manuscript. This work was supported by the European Research Council (ERC) under the European Union’s Horizon 2020 research and innovation programme (grant agreement No. 818808) and by Grant No. 2017239 from the United States-Israel Binational Science Foundation (BSF).

## Supplementary Information

**Figure S1.**
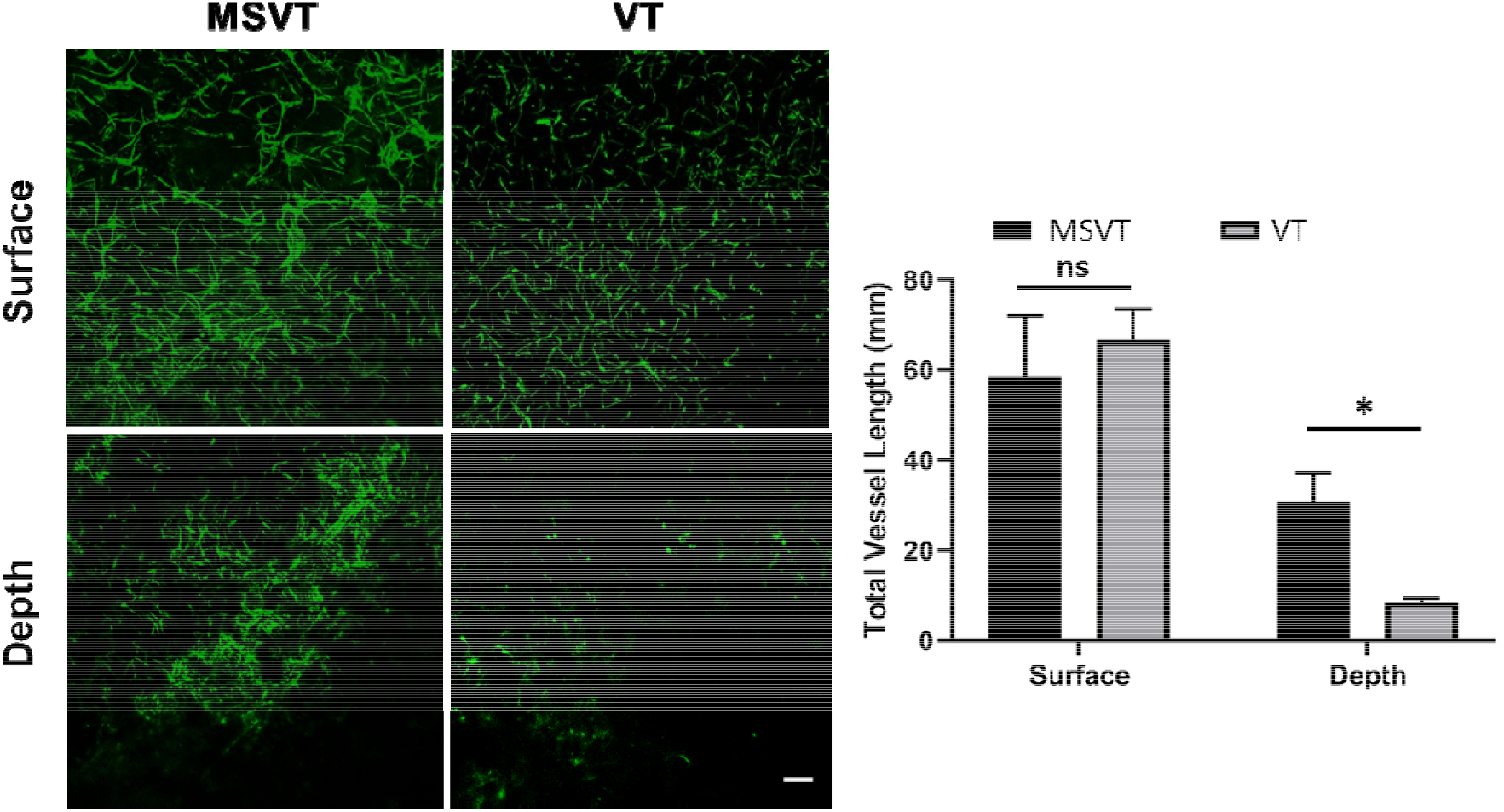
The impact of macro-channels on vascular depth distribution. (A) Representative images of construct-surface and in-depth MSVT and VT vasculature development, as determined by imaging HAMEC ZS green. (Scale bar: 200μm) (B) Quantification of total vessel length on the construct surface and in the construct depth (500μm) for MSVT and VT. Data are presented as means ± SEM. n = 4 (*P < 0.05).

**Figure S2.**
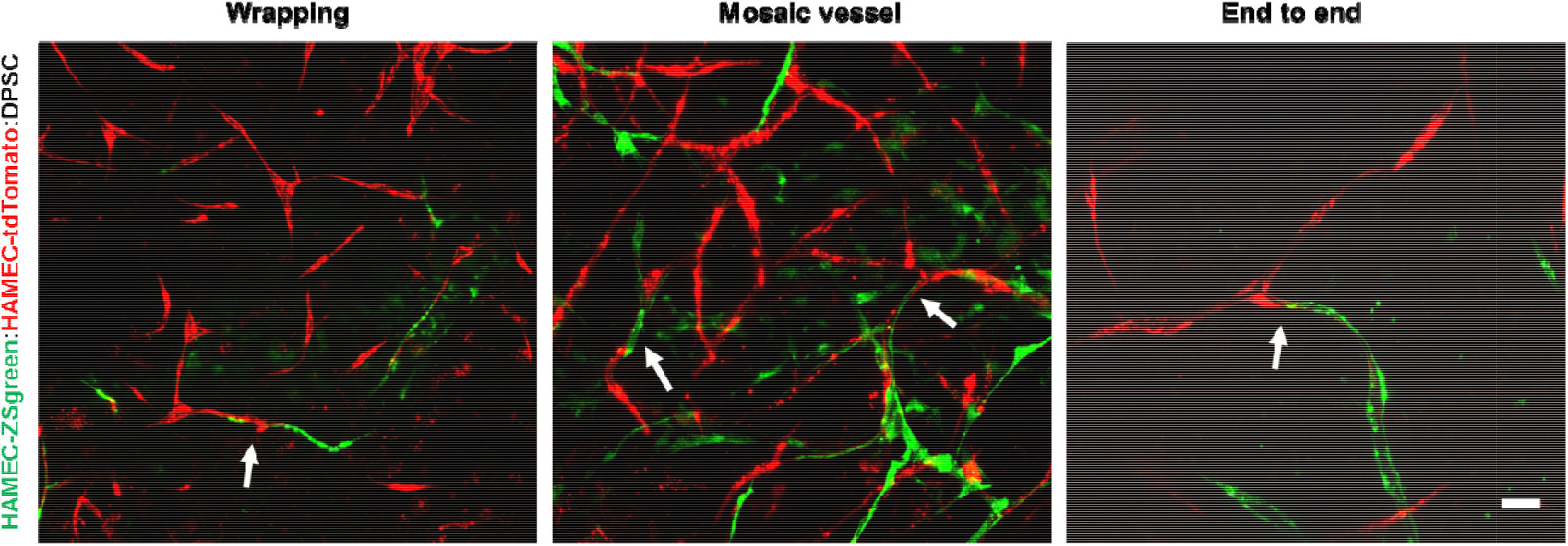
In vitro multi-scale vasculature interconnections. During in vitro cultivation of MVST, three different types of interconnections were observed between macrovessel sprouts and the surrounding self-assembled microvasculature, including: vessel wrapping (left), mosaic vessels (middle) and end to-end vessel anastomoses (right). (Scale bar: 50μm)

**Figure S3.**
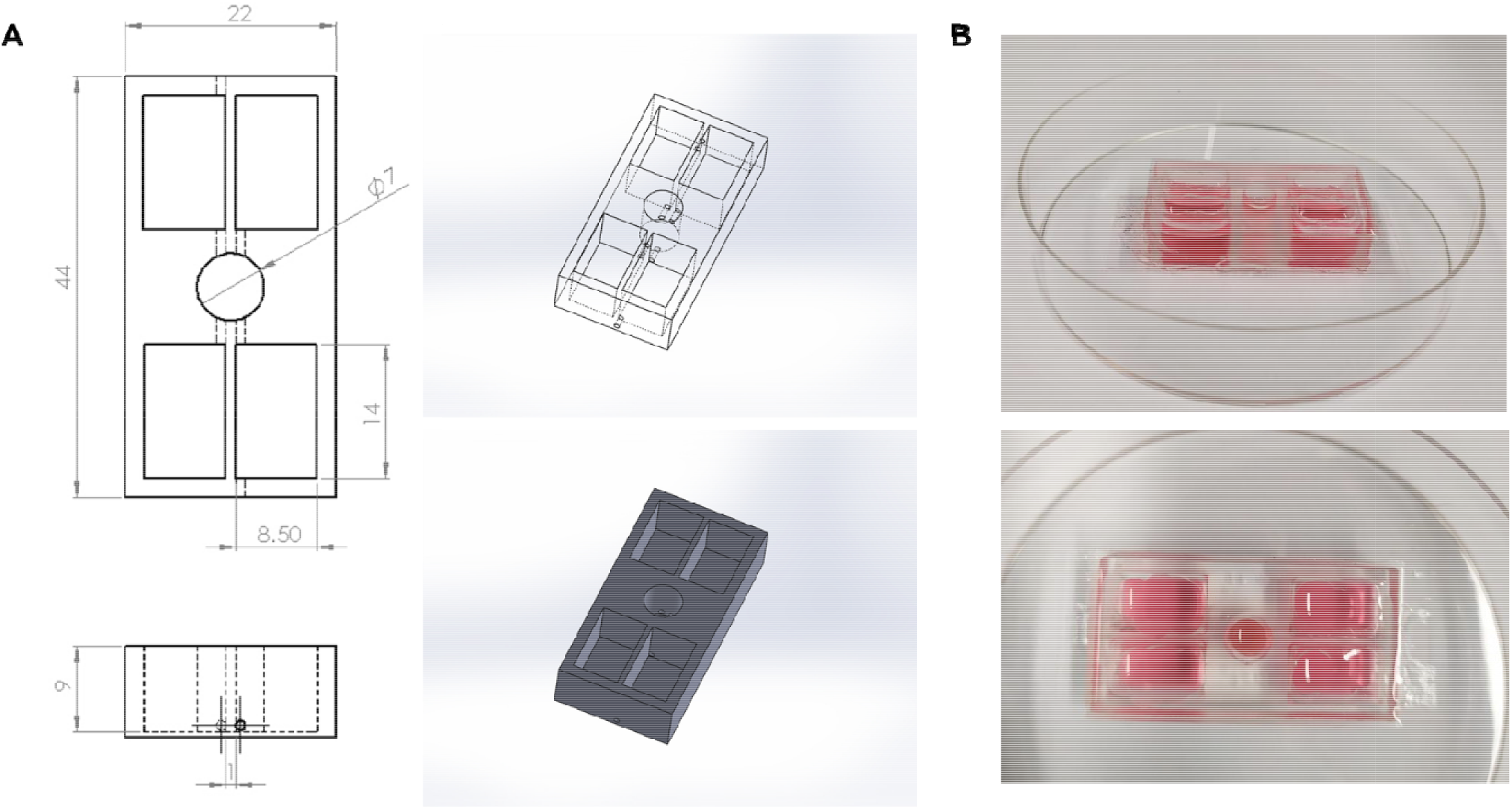
In-house-constructed flow bioreactor for MSVT cultivation. (A) Pumpless flow bioreactor design, manufacturing drawing and 3D CAD model. (B) PDMS flow bioreactor images.

**Figure S4.**
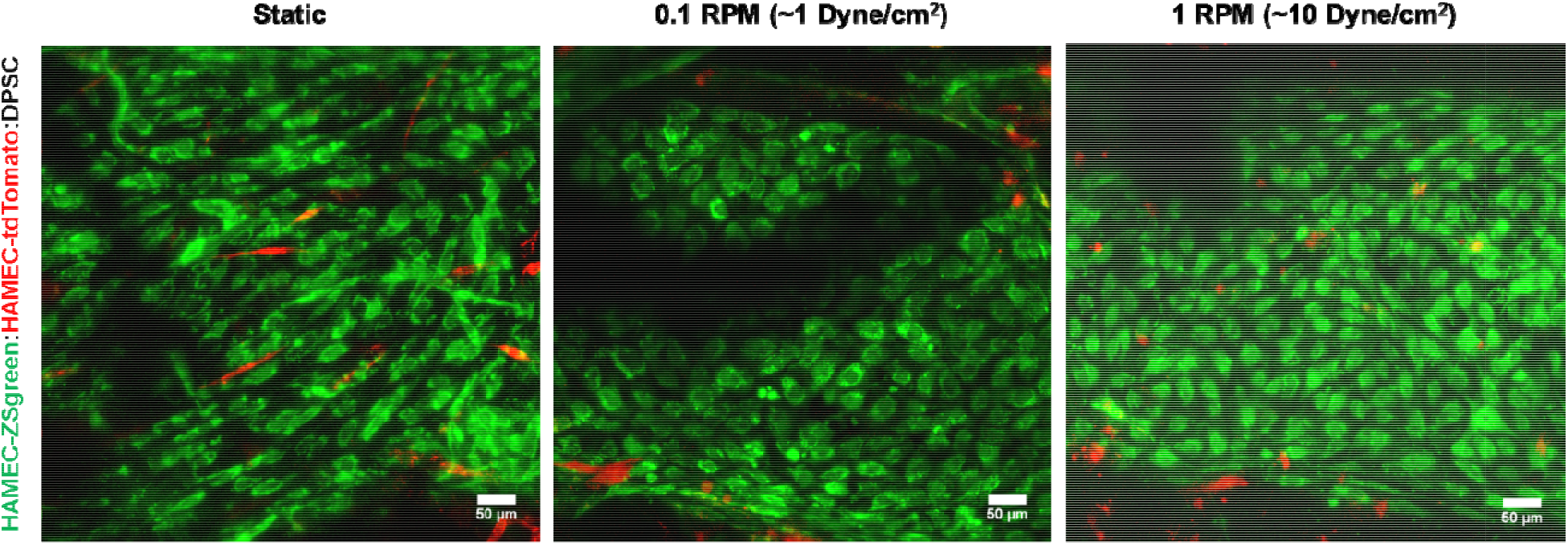
The impact of different shear stress levels on endothelium structure. Representative images of an engineered macro vessel endothelium (lined with HAMEC ZSgreen). Both low (middle) and high (right) flow rates induced a mature endothelium morphology, with a low aspect ratio and round cell shape (for early flow profile), while under static conditions (left), cells were characterized by a more elongated shape and high aspect ratio. (Scale bar: 50μm)

**Figure S5.**
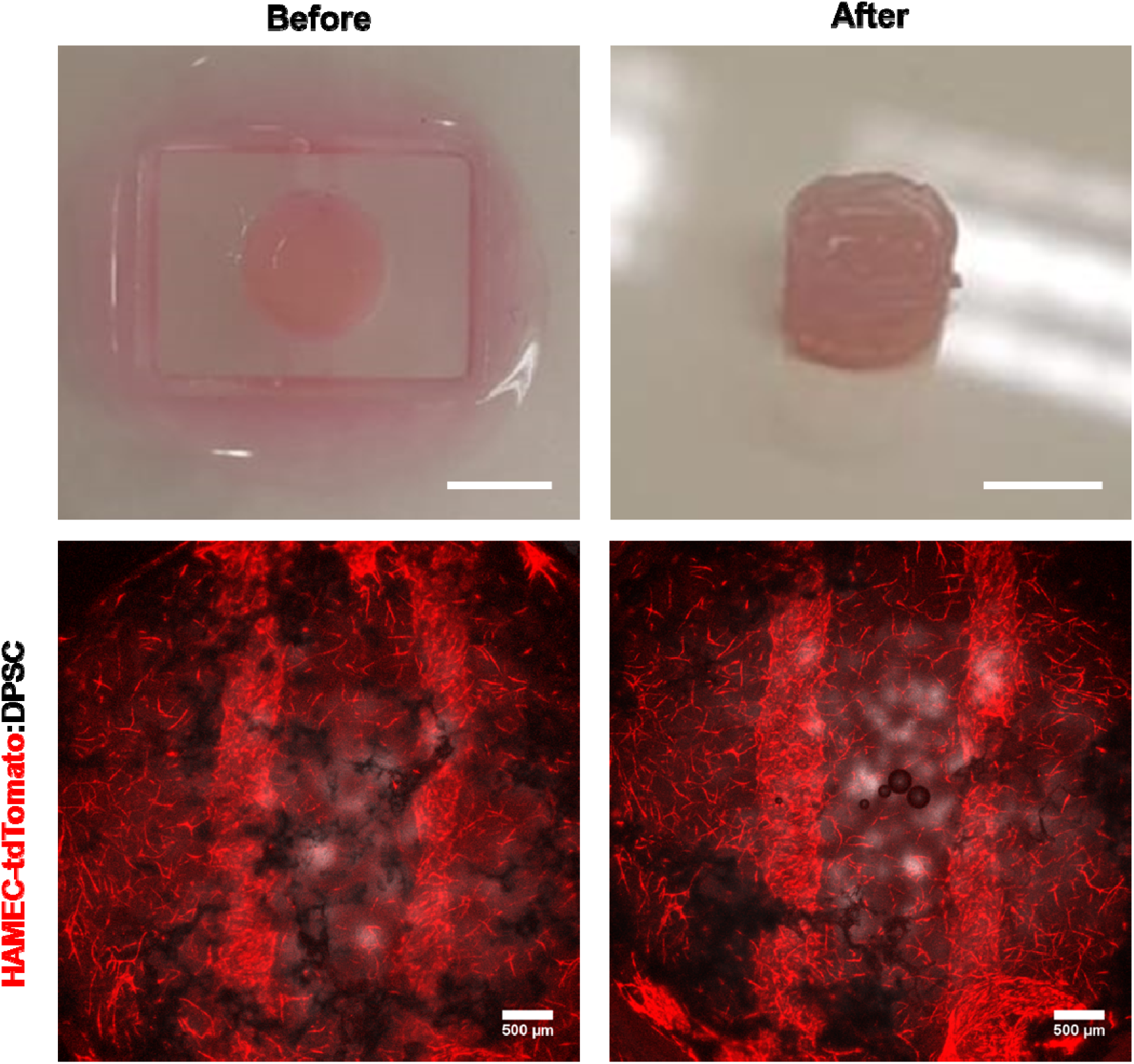
Vascular structure maintained stable after scaffold release. Low magnification and maximum intensity projection images of MSVT before and after release from the PDMS mold pre-implantation. The maintenance of the multi-scale vascular structure can be observed. (Scale bar: 5mm (upper); 500μm(lower))

**Figure S6.**
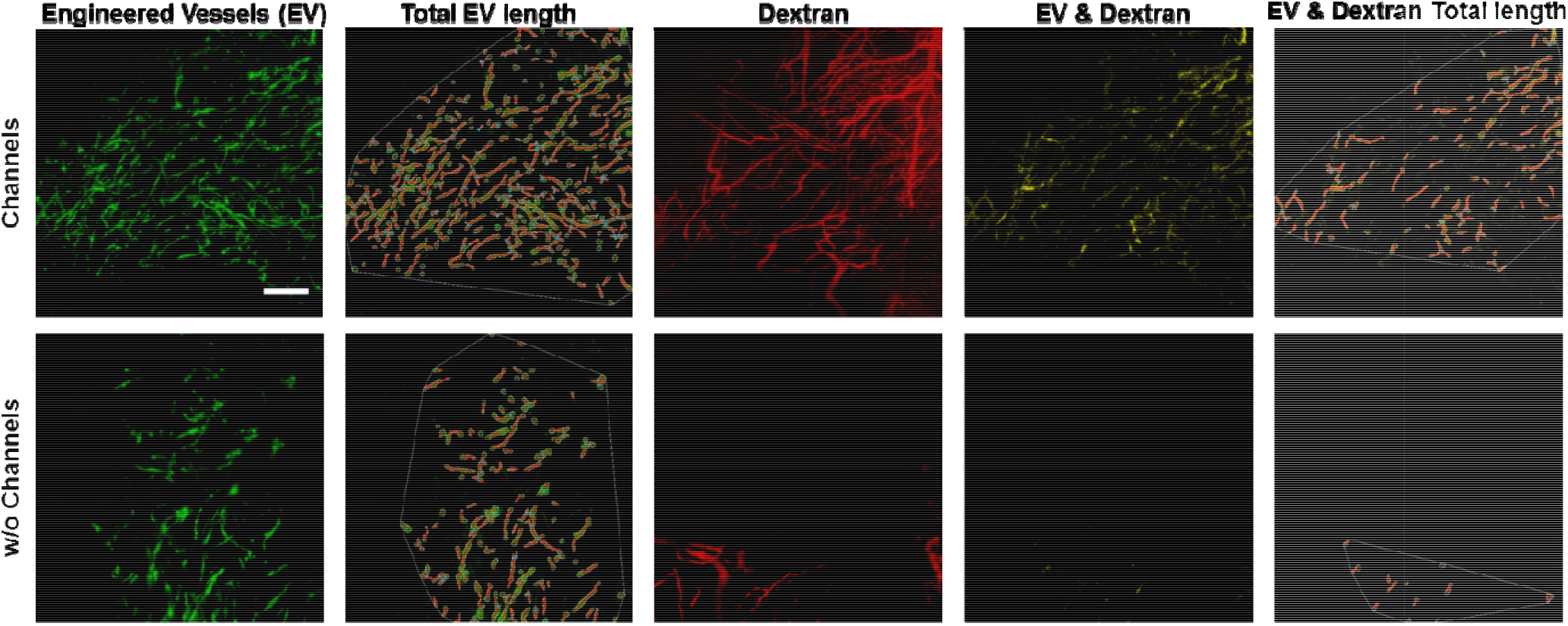
Quantification of perfused engineered vessel length using AngioTool. Total engineered vessel (EV) length was measured, followed by EV and dextran co-localization and length quantification. The perfused EV rate determined by dividing perfused EV length by total EV length. (Scale bar: 200μm)

## References

1. A. Lee, A. R. Hudson, D. J. Shiwarski, J. W. Tashman, T. J. Hinton, S. Yerneni, J. M. Bliley, P. G. Campbell, A. W. Feinberg, 3D bioprinting of collagen to rebuild components of the human heart. Science (80-.). 365 (2019).

2. N. Noor, A. Shapira, R. Edri, I. Gal, L. Wertheim, T. Dvir, 3D Printing of Personalized Thick and Perfusable Cardiac Patches and Hearts. Adv. Sci. 6, 1900344 (2019).

3. M. A. Skylar-Scott, S. G. M. Uzel, L. L. Nam, J. H. Ahrens, R. L. Truby, S. Damaraju, J. A. Lewis, Biomanufacturing of organ-specific tissues with high cellular density and embedded vascular channels. Sci. Adv. 5, eaaw2459 (2019).

4. J. S. Miller, K. R. Stevens, M. T. Yang, B. M. Baker, D.-H. T. Nguyen, D. M. Cohen, E. Toro, A. A. Chen, P. A. Galie, X. Yu, R. Chaturvedi, S. N. Bhatia, C. S. Chen, Rapid casting of patterned vascular networks for perfusable engineered three-dimensional tissues. Nat. Mater. 11, 768–774 (2012).

5. D. B. Kolesky, K. A. Homan, M. A. Skylar-Scott, J. A. Lewis, Three-dimensional bioprinting of thick vascularized tissues. Proc. Natl. Acad. Sci. 113, 3179–3184 (2016).

6. N. Y. C. Lin, K. A. Homan, S. S. Robinson, D. B. Kolesky, N. Duarte, A. Moisan, J. A. Lewis, Renal reabsorption in 3D vascularized proximal tubule models. Proc. Natl. Acad. Sci. U. S. A. 116, 5399–5404 (2019).

7. V. K. Lee, A. M. Lanzi, H. Ngo, S. S. Yoo, P. A. Vincent, G. Dai, Generation of multi-scale vascular network system within 3D hydrogel using 3D bio-printing technology. Cell. Mol. Bioeng. 7, 460–472 (2014).

8. A. Lesman, J. Koffler, R. Atlas, Y. J. Blinder, Z. Kam, S. Levenberg, Engineering vessel-like networks within multicellular fibrin-based constructs. Biomaterials 32, 7856–7869 (2011).

9. Y. J. Blinder, A. Freiman, N. Raindel, D. J. Mooney, S. Levenberg, Vasculogenic dynamics in 3D engineered tissue constructs. Sci. Rep. 5, 17840 (2016).

10. S. Levenberg, J. Rouwkema, M. Macdonald, E. S. Garfein, D. S. Kohane, D. C. Darland, R. Marini, C. A. van Blitterswijk, R. C. Mulligan, P. A. D’Amore, R. Langer, Engineering vascularized skeletal muscle tissue. Nat. Biotechnol. 23, 879–884 (2005).

11. S. Ben-Shaul, S. Landau, U. Merdler, S. Levenberg, Mature vessel networks in engineered tissue promote graft–host anastomosis and prevent graft thrombosis. Proc. Natl. Acad. Sci. U. S. A. 116 (2019).

12. D.-H. T. Nguyen, S. C. Stapleton, M. T. Yang, S. S. Cha, C. K. Choi, P. A. Galie, C. S. Chen, Biomimetic model to reconstitute angiogenic sprouting morphogenesis in vitro. Proc. Natl. Acad. Sci. 110, 6712–6717 (2013).

13. V. van Duinen, D. Zhu, C. Ramakers, A. J. van Zonneveld, P. Vulto, T. Hankemeier, Perfused 3D angiogenic sprouting in a high-throughput in vitro platform. Angiogenesis 22, 157–165 (2019).

14. T. Osaki, T. Kakegawa, T. Kageyama, J. Enomoto, T. Nittami, J. Fukuda, Acceleration of Vascular Sprouting from Fabricated Perfusable Vascular-Like Structures. PLoS One 10, e0123735 (2015).

15. X. Wang, D. T. T. Phan, A. Sobrino, S. C. George, C. C. W. Hughes, A. P. Lee, Engineering anastomosis between living capillary networks and endothelial cell-lined microfluidic channels. Lab Chip 16 (2016).

16. S. Kim, H. Lee, M. Chung, N. L. Jeon, Engineering of functional, perfusable 3D microvascular networks on a chip. Lab Chip 13, 1489–1500 (2013).

17. J. S. Jeon, S. Bersini, J. A. Whisler, M. B. Chen, G. Dubini, J. L. Charest, M. Moretti, R. D. Kamm, Generation of 3D functional microvascular networks with human mesenchymal stem cells in microfluidic systems. Integr. Biol. (United Kingdom) 6, 555–563 (2014).

18. M. A. Redd, N. Zeinstra, W. Qin, W. Wei, A. Martinson, Y. Wang, R. K. Wang, C. E. Murry, Y. Zheng, Patterned human microvascular grafts enable rapid vascularization and increase perfusion in infarcted rat hearts. Nat. Commun. 10, 1–14 (2019).

19. R. Sooppan, S. J. Paulsen, J. Han, A. H. Ta, P. Dinh, A. C. Gaffey, C. Venkataraman, A. Trubelja, G. Hung, J. S. Miller, P. Atluri,*In Vivo* Anastomosis and Perfusion of a Three-Dimensionally-Printed Construct Containing Microchannel Networks. Tissue Eng. Part C Methods 22, 1–7 (2016).

20. B. Zhang, M. Montgomery, M. D. Chamberlain, S. Ogawa, A. Korolj, A. Pahnke, L. A. Wells, S. Masse, J. Kim, L. Reis, A. Momen, S. S. Nunes, A. R. Wheeler, K. Nanthakumar, G. Keller, M. V. Sefton, M. Radisic, Biodegradable scaffold with built-in vasculature for organ-on-a-chip engineering and direct surgical anastomosis. Nat. Mater. 15, 669–678 (2016).

21. Y. S. J. Li, J. H. Haga, S. Chien, Molecular basis of the effects of shear stress on vascular endothelial cells. J. Biomech. 38, 1949–1971 (2005).

22. V. K. Lee, D. Y. Kim, H. Ngo, Y. Lee, L. Seo, S.-S. Yoo, P. A. Vincent, G. Dai, Creating Perfused Functional Vascular Channels Using 3D Bio-Printing Technology. Biomaterials 35, 8092–8102 (2014).

23. K. M. Gray, K. M. Stroka, Vascular endothelial cell mechanosensing: New insights gained from biomimetic microfluidic models. Semin. Cell Dev. Biol. 71, 106–117 (2017).

24. K. M. Chrobak, D. R. Potter, J. Tien, Formation of perfused, functional microvascular tubes in vitro. Microvasc. Res. 71, 185–196 (2006).

25. M. L. Moya, Y. H. Hsu, A. P. Lee, C. W. H. Christopher, S. C. George, In vitro perfused human capillary networks. Tissue Eng. - Part C Methods 19, 730–737 (2013).

26. A. Lesman, J. Koffler, R. Atlas, Y. J. Blinder, Z. Kam, S. Levenberg, Engineering vessel-like networks within multicellular fibrin-based constructs. Biomaterials 32, 7856–7869 (2011).

27. D. Rosenfeld, S. Landau, Y. Shandalov, N. Raindel, A. Freiman, E. Shor, Y. Blinder, H. H. Vandenburgh, D. J. Mooney, S. Levenberg, Morphogenesis of 3D vascular networks is regulated by tensile forces. Proc. Natl. Acad. Sci. U. S. A. 113, 3215–20 (2016).

28. Y. Shandalov, D. Egozi, A. Freiman, D. Rosenfeld, S. Levenberg, A method for constructing vascularized muscle flap. Methods 84, 70–5 (2015).

29. S. Levenberg, J. Rouwkema, M. Macdonald, E. S. Garfein, D. S. Kohane, D. C. Darland, R. Marini, C. A. van Blitterswijk, R. C. Mulligan, P. A. D’Amore, R. Langer, Engineering vascularized skeletal muscle tissue. Nat. Biotechnol. 23, 879–884 (2005).

30. A. A. Szklanny, L. Debbi, U. Merdler, D. Neale, A. Muñiz, B. Kaplan, S. Guo, J. Lahann, S. Levenberg, HighLThroughput Scaffold System for Studying the Effect of Local Geometry and Topology on the Development and Orientation of Sprouting Blood Vessels. Adv. Funct. Mater., 1901335 (2019).

31. W. Zhu, X. Qu, J. Zhu, X. Ma, S. Patel, J. Liu, P. Wang, C. S. E. Lai, M. Gou, Y. Xu, K. Zhang, S. Chen, Direct 3D bioprinting of prevascularized tissue constructs with complex microarchitecture. Biomaterials 124, 106–115 (2017).

32. C. Arakawa, C. Gunnarsson, C. Howard, M. Bernabeu, K. Phong, E. Yang, C. A. DeForest, J. D. Smith, Y. Zheng, Biophysical and biomolecular interactions of malaria-infected erythrocytes in engineered human capillaries. Sci. Adv. 6, eaay7243 (2020).

33. B. Grigoryan, S. J. Paulsen, D. C. Corbett, D. W. Sazer, C. L. Fortin, A. J. Zaita, P. T. Greenfield, N. J. Calafat, J. P. Gounley, A. H. Ta, F. Johansson, A. Randles, J. E. Rosenkrantz, J. D. Louis-Rosenberg, P. A. Galie, K. R. Stevens, J. S. Miller, Multivascular networks and functional intravascular topologies within biocompatible hydrogels. Science (80-.). 364 (2019).

34. O. Kérourédan, J. M. Bourget, M. Rémy, S. Crauste-Manciet, J. Kalisky, S. Catros, N. B. Thébaud, R. Devillard, Micropatterning of endothelial cells to create a capillary-like network with defined architecture by laser-assisted bioprinting. J. Mater. Sci. Mater. Med. 30, 1–12 (2019).

35. J. D. Baranski, R. R. Chaturvedi, K. R. Stevens, J. Eyckmans, B. Carvalho, R. D. Solorzano, M. T. Yang, J. S. Miller, S. N. Bhatia, C. S. Chen, Geometric control of vascular networks to enhance engineered tissue integration and function. Proc. Natl. Acad. Sci. 110, 7586–7591 (2013).

36. L. L. Y. Chiu, M. Montgomery, Y. Liang, H. Liu, M. Radisic, Perfusable branching microvessel bed for vascularization of engineered tissues. Proc. Natl. Acad. Sci. U. S. A. 109, E3414–E3423 (2012).

37. G. Cheng, S. Liao, H. K. Wong, D. A. Lacorre, E. Di Tomaso, P. Au, D. Fukumura, R. K. Jain, L. L. Munn, Engineered blood vessel networks connect to host vasculature via wrapping-and-tapping anastomosis. Blood 118, 4740–4749 (2011).

38. P. A. Galie, D. H. T. Nguyen, C. K. Choi, D. M. Cohen, P. A. Janmey, C. S. Chen, Fluid shear stress threshold regulates angiogenic sprouting. Proc. Natl. Acad. Sci. U. S. A. 111, 7968–7973 (2014).

39. D. C. Fernandes, L. S. Araujo, F. R. M. Laurindo, L. Y. Tanaka, Hemodynamic Forces in the Endothelium: From Mechanotransduction to Implications on Development of Atherosclerosis (2018) https:/doi.org/10.1016/B978-0-12-812348-5.00007-6 (April 22, 2020).

40. A. Reinitz, J. Destefano, M. Ye, A. D. Wong, P. C. Searson, Human brain microvascular endothelial cells resist elongation due to shear stress (2015) https:/doi.org/10.1016/j.mvr.2015.02.008 (April 22, 2020).

41. M. Sato, N. Kataoka, N. Ohshima, “Response of Vascular Endothelial Cells to Flow Shear Stress: Phenomenological Aspect” in Biomechanics, (Springer Japan, 1996), pp. 3–27.

42. L. Atia, D. Bi, Y. Sharma, J. A. Mitchel, B. Gweon, S. A. Koehler, S. J. Decamp, B. Lan, J. H. Kim, R. Hirsch, A. F. Pegoraro, K. H. Lee, J. R. Starr, D. A. Weitz, A. C. Martin, J. A. Park, J. P. Butler, J. J. Fredberg, Geometric constraints during epithelial jamming. Nat. Phys. 14, 613–620 (2018).

43. J. W. Song, L. L. Munn, Fluid forces control endothelial sprouting. Proc. Natl. Acad. Sci. U. S. A. 108, 15342–15347 (2011).

44. J. Koffler, K. Kaufman-Francis, Y. Shandalov, D. Egozi, D. Amiad Pavlov, A. Landesberg, S. Levenberg, A. P. Daria, A. Landesberg, S. Levenberg, Improved vascular organization enhances functional integration of engineered skeletal muscle grafts. Proc. Natl. Acad. Sci. 108, 14789–14794 (2011).

45. A. Lesman, M. Habib, O. Caspi, A. Gepstein, G. Arbel, S. Levenberg, L. Gepstein, Transplantation of a tissue-engineered human vascularized cardiac muscle. Tissue Eng. - Part A 16, 115–125 (2010).

